# Geographic cline analysis as a tool for studying genome-wide variation: a case study of pollinator-mediated divergence in a monkeyflower

**DOI:** 10.1101/036954

**Authors:** Sean Stankowski, James M. Sobel, Matthew A. Streisfeld

## Abstract

A major goal of speciation research is to reveal the genomic signatures that accompany the speciation process. Genome scans are routinely used to explore genome-wide variation and identify highly differentiated loci that may contribute to ecological divergence, but they do not incorporate spatial, phenotypic, or environmental data that might enhance outlier detection. Geographic cline analysis provides a potential framework for integrating diverse forms of data in a spatially-explicit framework, but it has not been used to study genome-wide patterns of divergence. Aided by a first-draft genome assembly, we combine an *F_CT_* scan and geographic cline analysis to characterize patterns of genome-wide divergence between divergent pollination ecotypes of *Mimulus aurantiacus*. F_CT_ analysis of 58,872 SNPs generated via RADseq revealed little ecotypic differentiation (mean *F_CT_* = 0.041), though a small number of loci were moderately to highly diverged. Consistent with our previous results from the gene *MaMyb2*, which contributes to differences in flower color, 130 loci have cline shapes that recapitulate the spatial pattern of trait divergence, suggesting that they reside in or near the genomic regions that contribute to pollinator isolation. In the narrow hybrid zone between the ecotypes, extensive admixture among individuals and low linkage disequlibrium between markers indicate that outlier loci are scattered throughout the genome, rather than being restricted to one or a few regions. In addition to revealing the genomic consequences of ecological divergence in this system, we discuss how geographic cline analysis is a powerful but under-utilized framework for studying genome-wide patterns of divergence.

## Introduction

The last decade has seen a resurgence of interest in the role that ecological differences play in the origin of new species (Rundle and Nosil 2005; Mallet 2008; Schluter et al. 2009; Sobel et al. 2010; Nosil 2012). Despite long-standing debate, most researchers have now embraced the notion that ecologically-based divergent selection can generate barriers to gene flow, especially early in speciation when other forms of isolation are absent (Nosil 2012). Moreover, recent efforts have advanced the field of speciation research beyond discussions of the nature of species or the plausibility of different geographic modes of speciation, towards a mechanistic understanding of the different factors that contribute to the evolution of reproductive isolation (Butlin et al. 2008; Mallet 2008; Sobel et al. 2010; Nosil et al. 2012; Butlin et al. 2012; Seehausen et al. 2014). Broad access to high-throughput sequencing technologies has enabled key questions about speciation to be placed in a genomic context (Hohenlohe et al. 2010; Seehausen et al. 2014), and many methods commonly employed in population genetic studies have been applied to genome-wide data (e.g. Hohenlohe et al. 2010).

A particularly active area of research surrounds the patterns of genome-wide divergence that accompany the speciation process (Butlin et al. 2012; Nosil 2012; Seehausen et al. 2014). Outlier scans that estimate population genetic statistics at thousands to millions of loci have revealed highly heterogeneous patterns of genomewide differentiation across a diverse array of taxa (Turner et al. 2005; Harr 2006; Hohenlohe et al. 2010; Martin et al. 2013; Renaut et al. 2013; Sorria-Carrasco et al. 2014; Twyford and Freidman 2015; Roesti et al. 2015; Burri et al. 2015). In many cases, only a small fraction of loci are highly diverged between ecotypes, while the majority of the genome shows relatively low differentiation (Seehausen et al. 2014). While it is now clear that the interpretation of this pattern is not as straightforward as once thought (Ralph and Coop 2010; Cruickshank and Hahn 2014; Burri et al. 2015), a common explanation is that these ‘oulier loci’ are associated with genomic regions that contribute to divergent selection and the barrier to gene flow between populations (Seehausen et al. 2014).

Given the relative ease with which genome-wide variation can be assayed, it is not surprising that genome scans have become a common first step in identifying candidate genomic regions that are associated with ecological divergence and speciation (Sessehausen et al. 2014). However, traditional genome scans have a number of limitations that are likely to reduce their efficacy in identifying ecologically important loci. For example, in conventional scans, there is no way to integrate detailed spatial patterns of phenotypic or environmental variation into the analysis. Rather, individuals are assigned *a priori* to groups based on one or more phenotypic or ecological criteria. However, morphological and ecological characteristics are often distributed continuously across geographic gradients (Endler 1977). Thus, any categorization into discrete groups may fail to capture the important variation of interest. Consequently, genome-wide analyses should ideally incorporate these diverse data in a spatially explicit framework. An additional limitation of most genome scans is that they do not take advantage of natural zones of admixture between divergently adapted populations. However, when they are present, hybrid zones can provide unique opportunities to study patterns of segregation and recombination among loci, which may reveal details about the genomic architecture of local adaptation (Barton and Gale 1993; Jiggins and Mallet 2000). For example, patterns of admixture and linkage disequilibrium in hybrid zones can establish whether highly diverged loci are broadly distributed throughout the genome or restricted to one or a few genomic regions.

Geographic cline analysis provides a potentially powerful framework for integrating multiple forms of data into studies of genome-wide variation. Cline analysis involves fitting cline models to allele frequency and/or quantitative data (e.g., phenotypic and environmental data), and has long been used as a tool for inferring the relative strengths of selection acting among loci (Barton & Hewitt 1985; Szymura and Barton 1986; Barton & Gale 1993; Gale et al. 2008). Despite the power of this approach for estimating an array of parameters, it has never been used as a tool to characterize patterns of genome-wide variation. One possible impediment is that the existing theoretical framework developed in the 1980s and 90s requires just a handful of differentially fixed (or nearly diagnostic), unlinked loci (Barton & Hewitt 1985; Szymura & Barton 1986; Barton & Gale 1993). Indeed, recent studies have often continued the tradition of removing markers with substantial allele sharing from datasets prior to analysis (e.g. Larson et al. 2014; Baldassare et al 2014; Lafontaine et al. 2015). However, adaptive divergence may commonly result in relatively modest divergence of loci between populations (Barrett & Schluter 2007). For example, genetic markers that are linked to causal regions may show lower divergence and may not emerge as candidates in conventional selection scans, but cline analysis should be able to detect spatial gradients in allele frequency, even if the differences are modest. Moreover, because quantitative data can be included in this framework (Bridle et al. 2001; Stankowski 2013), the shapes of molecular marker clines can be directly compared to clines in phenotypic traits or environmental variables, providing further support for their association with local adaption.

In this study, we use an outlier scan and cline analysis as we begin to characterize the pattern of genome-wide divergence between pollination ecotypes of the perennial shrub, *Mimulus aurantiacus*. In San Diego County, California, there is a sharp geographic transition between red- and yellow-flowered ecotypes of *M. aurantiacus* (Streisfeld and Kohn, 2005). These ecotypes are extremely closely related to each other, with the western, red ecotype having evolved recently from an ancestral yellow-flowered form that is currently distributed in the east (Stankowski and Streisfeld 2015). Despite their recent shared evolutionary history, the ecotypes show striking phenotypic differences in their flowers (Fig. 1; Waayers, 1996; Tulig, 2000; Streisfeld & Kohn 2005). In addition to pronounced variation in flower color, the ecotypes differ extensively in floral size, shape, and reproductive organ placement, which led Grant (1981, 1993ab) to hypothesize that divergence was driven primarily by selection to maximize visitation and pollen transfer by alternate pollinators.

**Figure 1.**
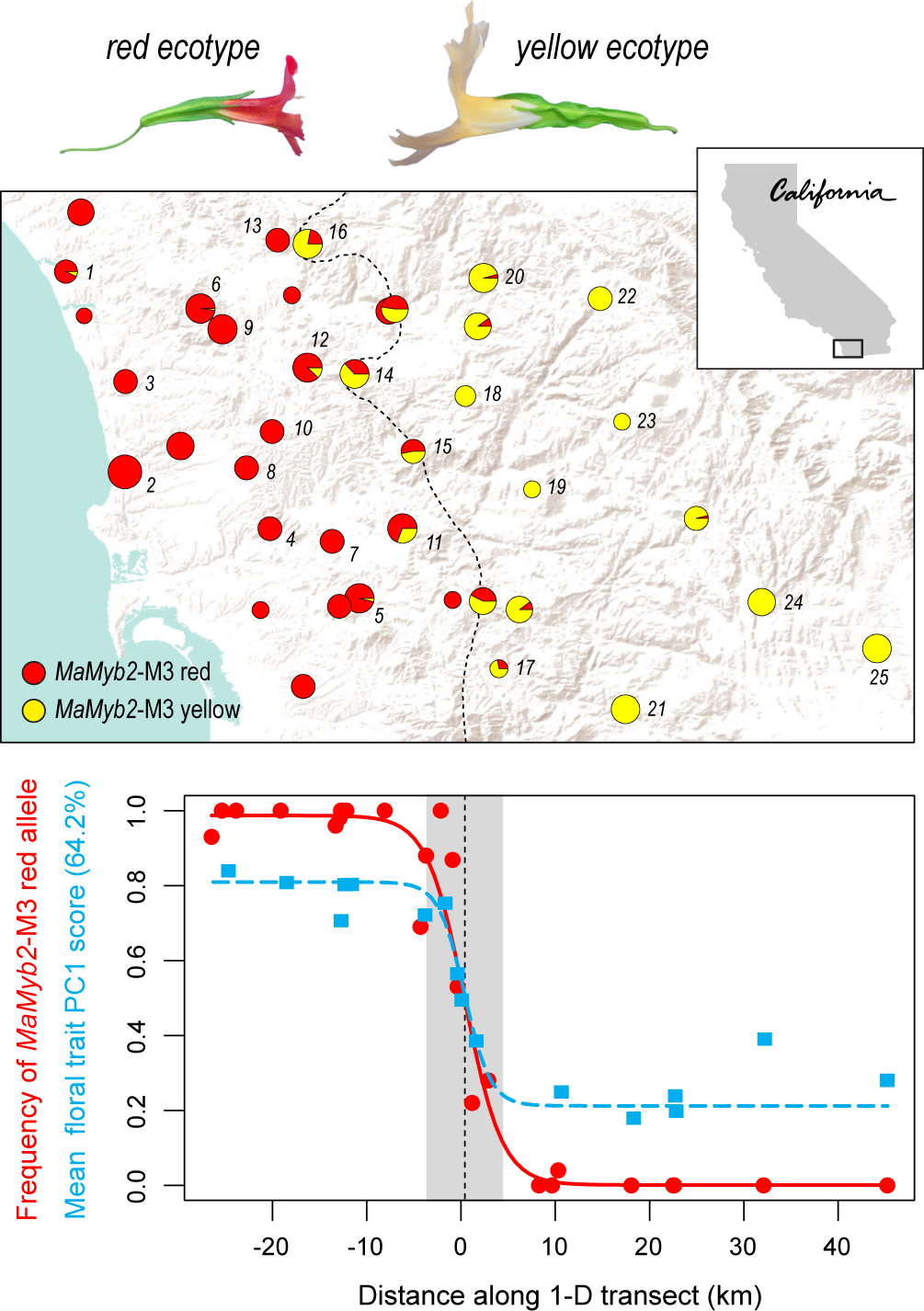
Sampling locations and clinal variation in MaMyb2-M3 and floral traits between the red and yellow ecotypes. The pie charts show the frequencies of *red* and *yellow* alleles at the *MaMyb2*-M3 marker across 39 locations. The dashed line is the contour where the alternative alleles are predicted to be at a frequency of 0.5. The 25 numbered sites are the locations selected for RADseq. b) One-dimensional (1-D) clines in *MaMyb2*-M3 allele frequency and mean floral trait PC1 score between the ecotypes. The allele frequency cline is based on allele frequencies from the 25 focus populations, while the floral trait PC1 cline (six phenotypic traits) is based on data from 16 locations (see Stankowski et al. 2015 for details)

Recent studies indicate that these phenotypic differences are maintained by divergent selection acting on floral traits despite ongoing gene flow. First, hummingbird and hawkmoth pollinators demonstrate opposing preferences and constancy for flowers of the red and yellow ecotypes, respectively, which generates strong but incomplete pollinator isolation between them (Streisfeld and Kohn 2007; Handelman and Kohn 2012; Sobel and Streisfeld 2015). A cis-regulatory mutation in the gene *MaMyb2* is primarily responsible for the transition from yellow to red flowers, and patterns of molecular variation in the gene show clear evidence for recent divergent selection (Streisfeld et al. 2013; Stankowski and Streisfeld 2015). In addition, six floral traits show sharp geographic clines across San Diego County (Stankowski et al. 2015). These clines are all positioned in the same geographic location, suggesting that the traits are differentiated due to a common selective agent. Therefore, we expect the loci underlying the floral traits to recapitulate the spatial position of trait divergence seen across San Diego County (Stankowski et al. 2015).

In contrast with the sharp transition in floral traits and the steep cline at *MaMyb2*, a recent study of more than 5000 single nucleotide polymorphisms (SNPs) revealed a gradual spatial pattern of divergence across the region, and patterns of floral trait variation in the hybrid zone are consistent with extensive admixture (Stankowski et al. 2015). These data support a long history of gene flow between the ecotypes and indicate that selection is responsible for maintaining floral trait differences. Further, endogenous post-mating barriers are effectively absent between the ecotypes, indicating that gene flow is not impeded by intrinsic hybrid unfitness (Sobel & Streisfeld 2015)

Here, we generated an initial draft genome assembly for *M. aurantiacus* and identified more than 50,000 SNP markers using RAD sequencing. After revealing limited genome-wide divergence between the ecotypes, we then fit a geographic cline model to each of the 427 most differentiated markers. To our knowledge, clines have never been fit to this many loci before. We find that estimates of cline shape and position outperform measures of allele frequency differentiation (*F_CT_*) in detecting loci that are associated with local adaptation. Indeed, we identified 130 loci whose cline shapes recapitulate the shape of the cline in floral trait divergence between the ecotypes. Patterns of admixture and linkage disequilibrium in the hybrid zone suggest that these loci are not restricted to one or a few genomic regions, suggesting that the loci contributing to local adaptation are broadly distributed throughout the genome. We end with a discussion of the features that make geographic cline analysis a powerful, but under-utilized approach for studying genome-wide variation, and highlight future directions for the further integration of cline analysis into the field of speciation genomics.

## Methods

### Genome sequencing and assembly

We sequenced and assembled a draft genome for *M. aurantiacus* using Illumina-based shotgun sequencing. We used a protocol outlined in Sobel and Streisfeld (2015) to isolate total genomic DNA from a greenhouse-grown individual of the red-flowered ecotype (Site UCSD; Table S1). We generated a single sequencing library by sonic shearing 1 ug of DNA, selecting the 400 – 600 bp size fraction and annealing paired-end T- overhang adapters to the repaired fragment ends (see supplement for a detailed protocol). After PCR enrichment of the library, 100bp paired-end sequencing was carried out in a single lane on the Illumina HiSeq 2000 at the University of Oregon’s Genomics Core Facility.

Initial processing of the raw reads was accomplished using the *Stacks* pipeline v. 1.12 (Catchen et al. 2013). The *processs-hortreads* program was used with default settings to discard reads with uncalled or low quality bases. The program *kmer_filter* was used to remove rare and abundant sequences over a range of different kmer sizes and abundance thresholds. After removing rare kmers that appeared only once and abundant kmers that were present more than 150,000 times, we used a kmer size of 69 to generate the final draft assembly using the software package *Velvet* (Zerbino and Birney 2008). Contigs of a minimum size of 100bp were retained, and summary statistics were calculated with custom scripts. Finally, as an assessment of the completeness of the gene space in our assembly, we used the *CEGMA* pipeline (Parra et al. 2007) to estimate the proportion of a set of 248 core eukaryotic genes (CEGs) that were completely or partially assembled. The proportion of CEGs present in an assembly has been shown to be correlated with the total proportion of assembled gene space, and thus serves as a good predictor of assembly completeness (Parra et al. 2009).

### Samples, RAD sequencing methods, and F_CT_ analysis

We identified single-nucleotide polymorphisms (SNPs) by sequencing restriction site associated DNA tags (RAD-seq) generated from 298 individuals sampled from 25 locations across the range of both ecotypes and the hybrid zone (mean individuals per site =12; range 4 to 18) in San Diego County, California (locations 1 to 25 in Fig. 1; Table S1). Samples from 16 of these populations were sequenced as part of a previous study that examined phenotypic divergence and population structure between the ecotypes (Stankowski et al. 2015). DNA isolation and sequencing libraries were prepared from an additional nine populations, following the methods described in Etter et al (2011), Sobel and Streisfeld (2015), and Stankowski et al. (2015). The 25 sample locations in the current study included 11 sites within the range of the red-flowered ecotype, 8 sites within the range of the yellow-flowered ecotype, and 6 sites located in the narrow transition zone where hybrid phenotypes have been observed (Stankowski et al. 2015) (Table S1).

We processed the raw sequencing reads, identified SNPs, and called genotypes using the *Stacks* pipeline v. 1.29 (Catchen et al. 2013). Reads were filtered based on quality, and errors in the barcode sequence or RAD site were corrected using the *process-radtags* script in *Stacks*. Individual reads were aligned to the *M. aurantiacus* genome (described herein) using *Bowtie 2*, with the *very-sensitive* settings. We then identified SNPs using the *ref_map.pl* function of *Stacks*, with two identical raw reads required to create a stack and two mismatches allowed when processing the catalog. SNP identification and genotype calls were conducted using the maximum-likelihood model implemented in *Stacks*, with alpha set to 0.01 (Hohenlohe et al. 2010, 2012; Catchen et al. 2011). We performed several independent runs in *Stacks* using a range of parameters for stack building and genotype calling, and all provided qualitatively similar results. To include a SNP in the final dataset, we required it to be present in at least 90% of all individuals and in a minimum of 8 copies across the entire dataset (i.e. minor allele frequency > 0.015).

We performed a locus-by-locus Analysis of Molecular Variance (AMOVA) in *Arlequin* v 3.5 (Excoffier *et al*. 2005) to determine the extent and pattern of genome-wide divergence between the red and yellow ecotypes. After accounting for variation partitioned between populations within the ecotypes and within populations, we obtained the fixation index F_CT_ between the ecotypes for each SNP marker. We arbitrarily defined markers in the top 1% of the F_CT_ distribution as “outlier loci,” and used these in subsequent analyses.

### Calculation of one-dimensional transect

We used one-dimensional (1-D) cline analysis to explore spatial variation in allele frequencies for each outlier locus. Although the ecotypes are distributed over a broad two-dimensional landscape, the transition between them occurs in a primarily east-west direction. To allow 1-D clines to be fitted to our data, we collapsed the two-dimensional sampling locations onto a 1-D transect. We used empirical Bayesian kriging, a geostatistical interpolation method, to generate a prediction surface of geographic variation in allele frequencies at the *MaMyb2-M3* marker, which is tightly linked to the *cis*-regulatory mutation that is primarily responsible for the transition from yellow to red flowers in *M. aurantiacus* (Streisfeld et al. 2013; Stankowski and Streisfeld 2015). Allele frequency data for 30 sample sites have been used in previous studies to generate a 1-D transect (Streisfeld et al. 2013; Stankowski et al. 2015). In this study, we include allele frequency data from nine additional locations that were sampled in spring of 2014 and genotyped according to Streisfeld *et al*. (2013) (Table S1).

We generated the prediction surface in *ArcMap* v. 10.2 (Esri) (see supplement for details), and determined the position of the cline center in two dimensions by extracting the linear contour where the frequencies of both alleles are predicted to be equal (*i.e*., 0.5). This location was set to position “0”. We then obtained 1-D coordinates for each of the 25 focal populations by calculating their minimum straight-line distance from the two-dimensional cline center, resulting in sites to the west of the two-dimensional cline center having negative 1-D distance values and sites to the east having positive values.

### The cline model and cline fitting procedure

With the 1-D transect established, we fitted a cline model to the allele frequency data for each outlier SNP (top 1% of the *F_CT_* distribution) using maximum likelihood (ML). As SNPs located within 95 bp on either side of the same restriction enzyme cut site (*PstI*) show very similar patterns of divergence (see results), we only fitted a cline to one randomly selected SNP in each 190 bp RAD locus (RAD tags are sequenced 95 bp in each direction from each *PstI* cut site).

We used a *tanh* cline model described in Szymura and Barton (1986, 1991) to obtain 5 parameters that describe the spatial position, rate, and extent of allele frequency changes across the 1-D transect. These parameters, which are either estimated during the fit, or derived from the ML solution, are described in greater detail in Figure 2. The first parameter, Δ*P*, provides an estimate of the total change in the allele frequency difference across the transect. Like *F_CT_*, Δ*P* can range between 0 (no difference in allele frequency across the cline) and 1 (alternative alleles fixed in each tail). However, unlike *F_CT_* analysis, where the difference in allele frequency is estimated between discrete groups, Δ*P* is estimated from the tails of a continuous function. Thus, depending upon how the allele frequencies vary over space, estimates of *F_CT_* and Δ*P* may differ markedly from one another. The second and third parameters, *P*_max_ and *P_min_*, describe the frequency of allele *i* in the high and low tails of the cline, respectively, which allows us to explore the spatial pattern of allele frequency variation in more detail. For example, both alleles at a locus may be present at appreciable frequencies on one side of the cline, but one of the alleles could be fixed on the other side of the cline. In this case, the marker would show a moderate Δ*P*, but *P_max_* and *P_min_* indicate different levels of allele sharing on each side of the cline.

**Figure 2.**
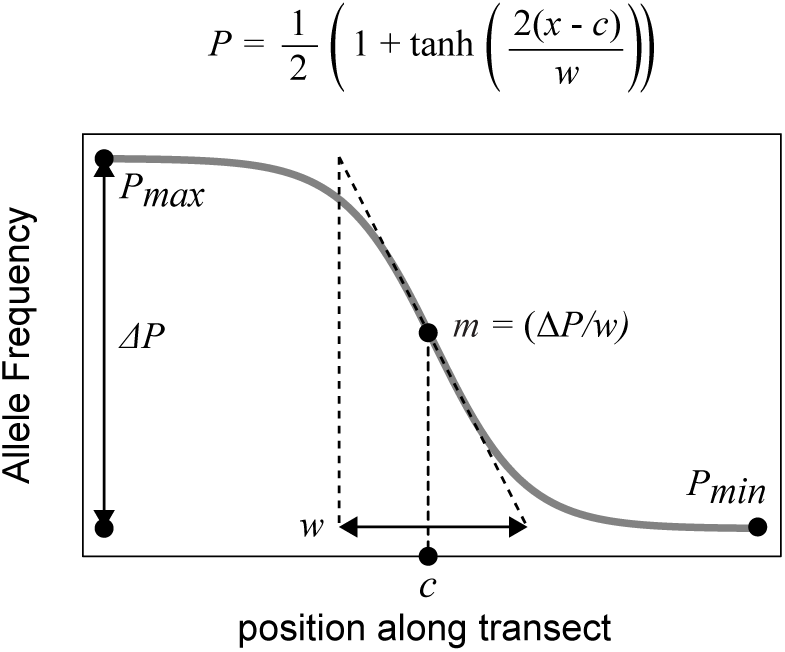
The sigmoid cline model. The hyperbolic *tanh* function enables us to estimate two parameters that describe cline shape, and derive four additional parameters from the ML solution. The estimated parameters are the cline centre (c), which is the geographic position of the maximum gradient of the cline function, and the cline width (w). The derived parameters are *P*_max_, the frequency of the focal allele in the high tail, *P*_min_, the frequency of the focal allele in the low tail, and Δ*P*, the total change in the allele frequency across the transect, calculated as *P*_max_ - *P*_min_. Finally, the cline slope, m, defined as the maximum gradient of the sigmoid function, is calculated as Δ*P*/*w*.

The fourth and fifth parameters are the cline center (*c*) and cline slope (*m*) (Fig. 2). Assuming that a cline is at equilibrium, and that alternative alleles are maintained in different areas due to selection across an ecological gradient, the cline centre indicates the position where the direction of selection acting on an allele changes (Endler 1977; Barton & Gale 1993; Krukk et al. 1999). The cline slope (*m*) indicates the rate of change in allele frequency at the maximum gradient of the function and provides a relative indication of the strength of selection acting on a locus, with cline slope increasing as the strength of selection increases. Traditionally, the comparison of cline width (*w*) has been used to infer variation in the strength of selection acting among loci (Barton and Hewitt 1985; Barton & Gale 1993). However, *w* is a function of Δ*P*, which means that for a given value of m, cline width decreases as Δ*P* decreases. In studies of genome-wide variation, we expect considerable variation in Δ*P*, which complicates comparisons of *w* among loci. Thus, when a set of markers shows considerable variation in allele sharing, cline slope has a more straightforward interpretation.

Cline fitting was conducted using the maximum likelihood framework implemented in *Analyse* v. 1.3 (Barton and Baird 1993). For each “outlier locus” (top 1% of the *F_CT_* distribution), and for the *MaMyb2-M3* marker, we fit a cline to the allele frequency data from the 25 sample sites, arbitrarily using the allele that was most common in the red ecotype as the focal allele. To ensure that the likelihood surface was thoroughly explored, we conducted two independent runs, each consisting of 10,000 iterations, with different starting parameters and random seeds. Each fit was visually inspected for quality. Because we were interested in identifying markers with cline shapes that were similar to floral traits, we re-fitted a cline to the floral trait data for the 16 populations published in Stankowski et al. (2015) using the new 1-D transect. Rather than fitting a cline to each trait, we conducted a Principal Components analysis on the trait data for each individual, and scaled the data between 0 and 1 as required by the software. We then calculated the mean PC1 score for each site, and fitted a onedimensional cline as described in Stankowski et al. (2015).

Given that linked loci often show similar patterns of divergence, we also tested whether markers in close genomic proximity have similar cline shapes. Because our genome assembly consists primarily of short scaffolds (see results), we were unable to perform a detailed analysis of cline parameters across large chromosomal regions. However, for each of the five cline parameters (*P_min_, P_max_*, Δ*P*, *c*, and *m*), we used a regression analysis to test for a relationship in the value of the parameter between all pairs of SNPs found on the same genome scaffold. We tested the significance of the relationship by comparing the observed *r*^2^ value to a null distribution of values generated using 1,000,000 random permutations of the data using custom scripts in *R*, as described in Stankowski et al. (2015).

### Associations between outlier loci in the hybrid zone

We have shown previously that floral trait associations are greatly reduced in the hybrid zone between the ecotypes, which is consistent with ongoing gene flow and recombination between the ecotypes (Stankowski et al. 2015). Such widespread hybridization provides us with an excellent opportunity to determine how the outlier loci are distributed throughout the genome. For example, if the loci are restricted to one or a few genomic regions, we expect the associations of alleles among loci to be maintained despite ongoing gene flow. However, if loci are spread throughout the genome, we expect the associations in the hybrid zone to be dramatically reduced.

We used two methods to assess the strength and pattern of the association among alleles in the outlier loci. First, we used *Structure v. 2.3.4* (Pritchard et al. 2000) to infer patterns of admixture across the outlier loci and the *MaMyb2-M3* marker, using the settings outlined in Stankowski et al. (2015). We then compared the distributions of hybrid index scores from individuals in the four hybrid populations examined in Stankowski et al (2015) (*n* = 61) relative to the pure red- and yellow-flowered ecotypes. Extensive admixture in the hybrid zone among these outlier loci would be reflected by a broad, unimodal distribution of hybrid index scores, suggesting that alleles at different loci are inherited independently of each other in hybrid individuals.

Second, we examined patterns of linkage disequilibrium (LD) among the outlier loci inside versus outside the hybrid zone. Significant reductions of LD in the hybrid zone would suggest that the markers are distributed primarily in different genomic regions. We first generated a theoretical expectation for LD in the absence of inter-ecotype gene flow by pooling individuals from pure red and yellow ecotype sites into a single ‘population.’ We calculated D’ separately in this pure ‘population’ and in samples in the hybrid zone between all pairs of outlier SNPs using the *R* package *LDheatmap* (Shin et al. 2006). We qualitatively evaluated whether the number of groups of loci that showed tight LD with one another in the hybrid zone was reduced relative to the pooled pure ‘population’. Specifically, we used the *heatmap2* function in the *R* package *gplots* to cluster sets of markers with similar *D*’ estimates in each of the pairwise matrices. We also tested for quantitative reductions in the mean estimates of D’ within the hybrid zone relative to the mean estimate obtained from our pooled population using a permutation test (100,000 permutations). However, because physical linkage and selection can influence associations between alleles among loci, we also tested for differences in mean LD inside and outside the hybrid zone for (*i*) pairs of markers located on different genome scaffolds, (*ii*) pairs of linked loci located on the same genome scaffold, and (*iii*) all pairwise combinations that included the *MaMyb2-M3* marker, which is in the divergently selected flower color locus *MaMyb2*.

## Results

### Draft Genome Assembly for Mimulus aurantiacus

We obtained 176,451,402 raw paired reads from a single lane of Illumina HiSeq PE100 sequencing of one red ecotype individual, which is equivalent to an average of 118x coverage of the estimated 297 Mbp genome size (Murovec and Bohanek 2013). The final draft assembly consisted of 23,129 scaffolds larger than 500bp and totaled 223.8 Mbp (74% of the estimated genome size). The N50 scaffold length was 31,153 bp, and the largest scaffold was 209,453 bp. *CEGMA* analysis identified partial or complete sequences for 240 of the 248 CEGs (97%), with 208 of them being completely assembled.

### F_CT_ analyszs reveals low genome-wzde dzvergence between the ecotypes

To explore genome-wide divergence between the ecotypes, we sequenced RAD tags from 298 individuals, aligned the reads to our draft assembly and identified 58,872 SNPs that met our filtering requirements. A locus-by-locus analysis of *F_CT_* for these markers revealed a highly skewed distribution of genetic divergence between the ecotypes, with most loci showing little or no differentiation (Fig. 3a). Estimates of *F_CT_* ranged from -0.062 to 0.853, with a mean inter-ecotype differentiation of 0.041 (s.d. 0.075). The top 1% of the *F_CT_* distribution (*n* = 589) spanned roughly half of the total range of values among RAD markers, with a minimum value of 0.358. These ‘outlier’ SNPs showed moderate differentiation between the ecotypes (mean 0.451 s.d. 0.084), but none of the markers were as highly differentiated as the *MaMyb2-M3* marker (*F_CT_* = 0.98).

**Figure 3.**
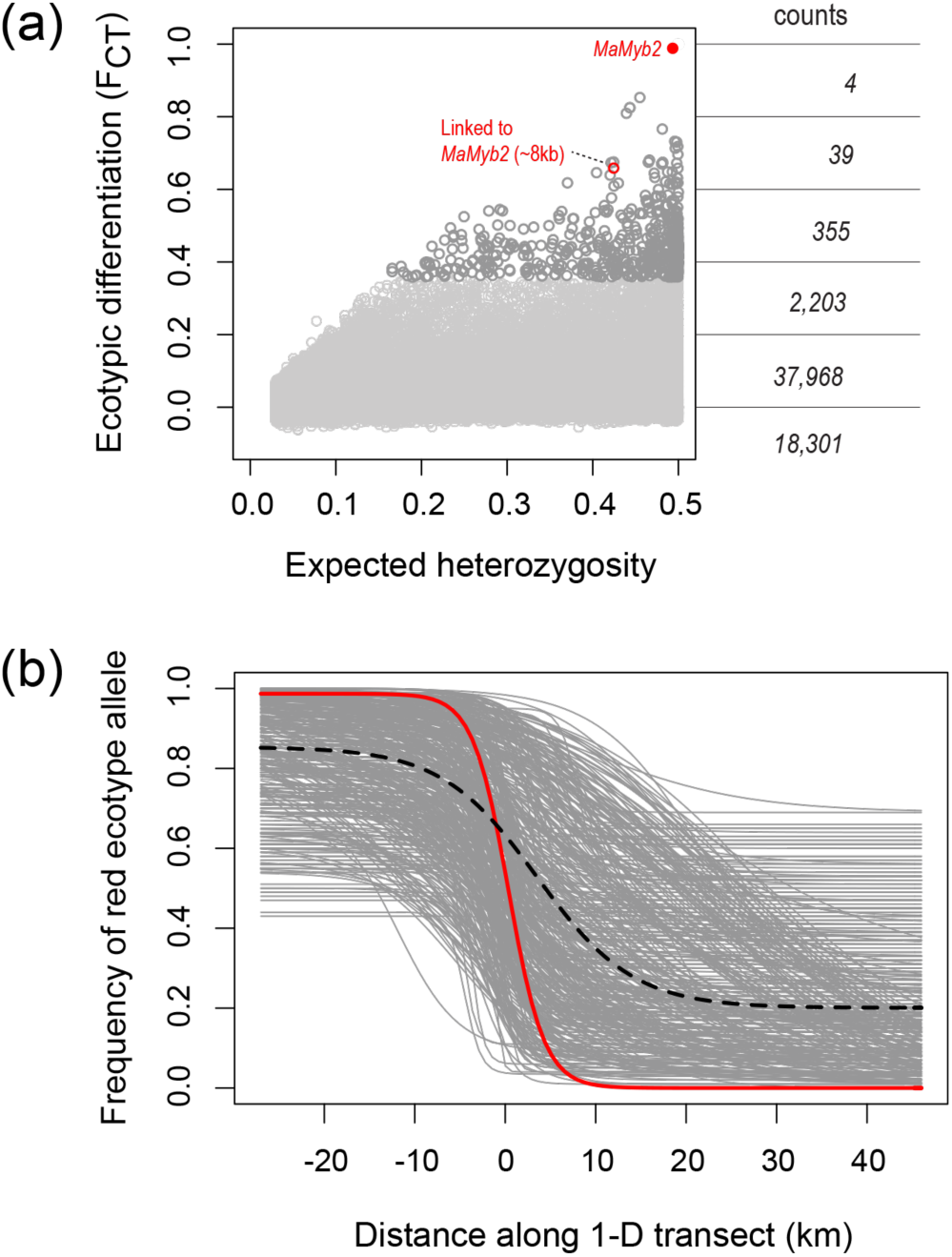
Differentiation of SNP markers between the red and yellow ecotypes. a) Plot of F_CT_ on expected heterozygosity for 58,872 RAD markers and the *MaMyb2-M3* marker. SNPs in the top 1 *%* of the F_CT_ distribution are colored dark gray (n = 589). The counts show the density of markers in six bins. The *MaMyb2-M3* marker and a SNP located approximately 8 kb from *MaMyb2* are highlighted. b) Geographic clines for the top 1% of the F_CT_ distribution, including only a single SNP marker per RAD cut site (n = 426). The red line shows the cline for the *MaMyb2-M3* marker; the dashed line is the average cline based on the mean parameter estimates across the 426 markers.

The 589 SNPs in the top 1% of the *F_CT_* distribution are located in 426 distinct RAD loci. We expected that SNPs found in the same locus would show very similar levels of inter-ecotype differentiation. Indeed, a pairwise regression of *F_CT_* among pairs of these 163 SNPs was highly significant and explained more than 90% of the variation in inter-ecotype differentiation (*r*^2^ = 0.91; permutation test *p* = 9.99 x10-7; Fig. S1). Therefore, we conducted further analyses based on data from a single randomly selected SNP from each of the 426 unique 190 bp RAD loci represented in the top 1% of the *F_CT_* distribution, as well as a marker in the *MaMyb2* gene (*MaMyb2-M3)*.

### Geographic cline analysis reveals extensive variation in the spatial pattern of divergence among loci

Consistent with previous results and the role of *MaMyb2* in flower color evolution (Streisfeld et al. 2013; Stankowski et al. 2015), the cline in the floral traits and *MaMyb2*- M3 marker had very similar shapes (Fig. 1). In contrast, we observed a diverse array of cline shapes for the 426 “outlier loci” (Fig. 3b). The average 1-D cline (calculated from the mean of all parameter estimates) was shallower than the cline for the *MaMyb2-M3* marker, both in terms of the total change in allele frequency across the 1-D transect (Δ*P*_mean_ = 0.66 vs. Δ*P*_uaMyb2_ = 0.99), and cline slope (*m_mean_* = 0.032 vs. *muaMyb2* = 0.125). The average cline center was shifted approximately 5 km to the east of the center for the floral trait cline.

Given the difficulty of drawing conclusions from visual analysis of so many clines, we examined the distributions of the parameters that describe cline shape (Fig. 4). As with *F_CT_*, the three parameters involving allele frequency change in the tails (Δ*P*, *P*_max_ and *P*_mim_), revealed considerable allele sharing between the ecotypes (Fig. 4). However, examination of the cline parameters provides additional information about the spatial patterns of this allele sharing that is not possible from estimates of *F*_ct_. Specifically, although 77% of markers were at or near fixation in at least one tail of the cline (40% had *P*_max_ > 0.90 in the left tail and 37% had *P*_min_ < 0.10 in the right tail; Fig. 4b), 75% of the 427 markers had a Δ*P* less than 0.8 (Fig. 4a). Thus, despite most markers being near fixation on one side of the cline, both alleles were often at appreciable frequency in the opposite tail.

Patterns of variation in the remaining two parameters, cline center (*c*) and slope (*m*) revealed a subset of loci with cline shapes that recapitulated the pattern and scale of floral trait divergence across San Diego County. First, despite broad variation in the estimates of *c* (range -13 km to 30 km), 60% of markers had ML estimates of cline center that coincided with the narrow phenotypic transition zone between the ecotypes, as defined by the width of the floral trait cline (*w* for the mean floral trait PC1 score = -3.5 km to 3.5 km; Fig. 4c). Cline slope varied more than 40-fold among loci, with estimates of *m* ranging from 0.008 to 0.344 (Fig. 4d). One-third of markers had slopes that were greater than the average for all 427 markers (mean *m* = 0.053), including a RAD locus located approximately 8 kb from the flower color gene *MaMyb2*. Fifty-six markers had slopes that were greater than the slope for the *MaMyb2-M3* marker (m = 0.125). Finally, we observed a striking relationship between cline center and cline slope (Fig. 4e), with the sharpest clines coinciding with the geographic position of the cline center in floral traits. Specifically, 130 marker clines had cline centers that coincided with the floral trait cline and whose slopes were elevated above the average for the 427 markers (*m*_mean_ = 0.053). This included both the *MaMyb2-M3* marker and the RAD marker located approximately 8 kb from *MaMyb2*.

**Figure 4.**
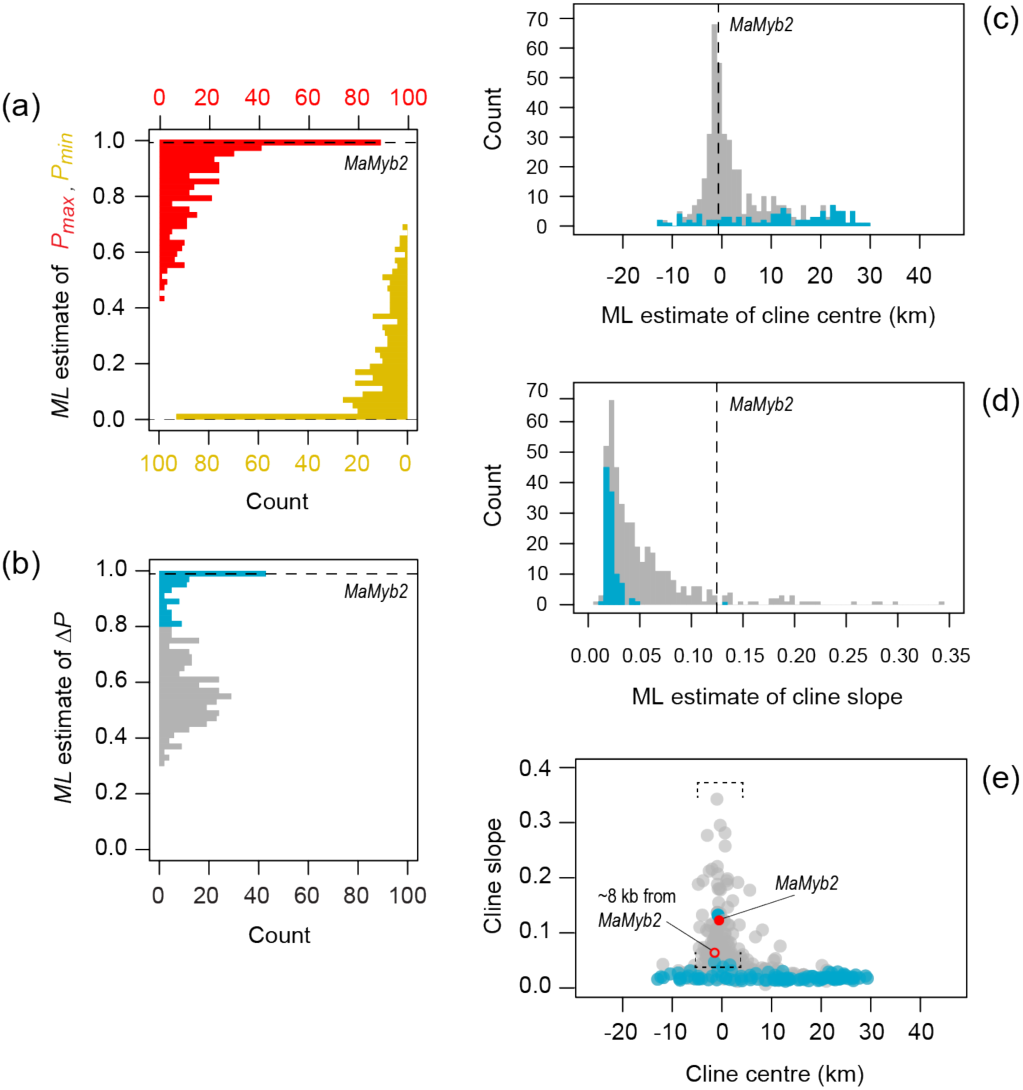
Distributions of ML cline parameters for the 426 RAD outlier loci. (a) Distributions of allele frequencies for *P*_max_ (red) and *P*_min_ (yellow) in the left and right tails of each cline. (b) Change in allele frequency across the cline (Δ*P*). (c) Distribution of cline center, (d) cline slope, and (e) the relationship between cline center and cline slope. The dashed black lines in plots a - d show the ML estimates for the *MaMyb2-M3* cline. In panels b-e, the blue bars and points are for markers where Δ*P* is > 0.8, and the gray bars and points show the distribution for markers where Δ*P* < 0.8. In panel e, points within the brackets have centers that coincide with the geographic transition in floral traits, and have cline slopes above the average for all 427 loci.

Curiously, our cline analysis also revealed that markers showing the largest differences in allele frequency in both tails (Δ*P* > 0.8) tended to have cline shapes that were discordant from the spatial pattern of trait divergence between the ecotypes (Fig. 4d). Specifically, these markers tended to have the shallowest slopes, and cline centers that were shifted to the east of the *MaMyb2-M3* cline (Fig. S2). Rather, the markers that showed cline shapes that recapitulated the spatial transition in the floral traits tended to show moderate differences in allele frequency across the transect (Δ*P* < 0.8) (Fig. S2).

### SNPs in close genomic proximity have similar cline shapes

Using our draft genome assembly, we tested whether SNPs in the same genomic regions had similar cline shapes. In support of this hypothesis, a regression including 97 pairs of loci from the 64 genomic scaffolds containing more that one outlier SNP (mean distance between SNPs = mean 9.7 kb s.d., 14.3 kb; max distance = 74.3 kb), explained 40% of the variation in Δ*P* (*r*^2^ = 0.400, *p* = 9.99 x 10^-7^), 51% and 37% of the variation in *P*_max_ and *P*_min_, respectively (*P*_max_: *r*^2^ = 0.509, *p* = 9.99 x 10^-7^; *P*_min_: *r*^2^ = 0.373, *p* = 9.99 x 10^-7^), 51% of the variation in cline center (*c*: *r*^2^ = 0.505, *p* = 9.99 x 10^-7^) and 35% of the variation in cline slope (*m*: *r*^2^ = 0.353; *p* = 4.99 x 10^-6^) (Fig. S3).

### F_CT_ is a poor predictor of variation in cline shape parameters

We tested for a relationship between ecotypic differentiation (*F_CT_*) and the ML estimates of each cline parameter obtained for the 427 markers. In general, *F_CT_* was a poor predictor of the parameters that describe cline shape. Although highly significant, only 3.4% of the variation in Δ*P* was explained by the estimates of *F_CT_*(*r*^2^ = 0.0346, *p* < 0.0001). For the other cline parameters, the relationships were even weaker. Estimates of *F_CT_* explained only 1.4 percent of the variation in cline center (*r*^2^ = 0.0137, *p* = 0.015), and 1.9% of the variation in cline slope (*r*^2^ = 0.0193, *p* = 0.0041).

### Associations among outlier loci are reduced in the face of gene flow

The observed patterns of admixture and linkage disequilibrium in the hybrid zone suggest these outlier loci are scattered broadly throughout the genome. Based on admixture scores, individuals from pure red and yellow flowered sample sites were generally assigned into alternative clusters with high probability (Fig. 5a). A few individuals were clear outliers in each distribution, suggesting they were either hybrids or pure individuals of the alternative ecotype. In contrast, the distribution of hybrid index scores in the hybrid zone spanned nearly the full range of values and had a roughly unimodal shape centered intermediate of the pure ecotypes. This extensive admixture indicates that the alternative alleles among these outlier loci are often inherited independently of one another in hybrid offspring.

**Figure 5.**
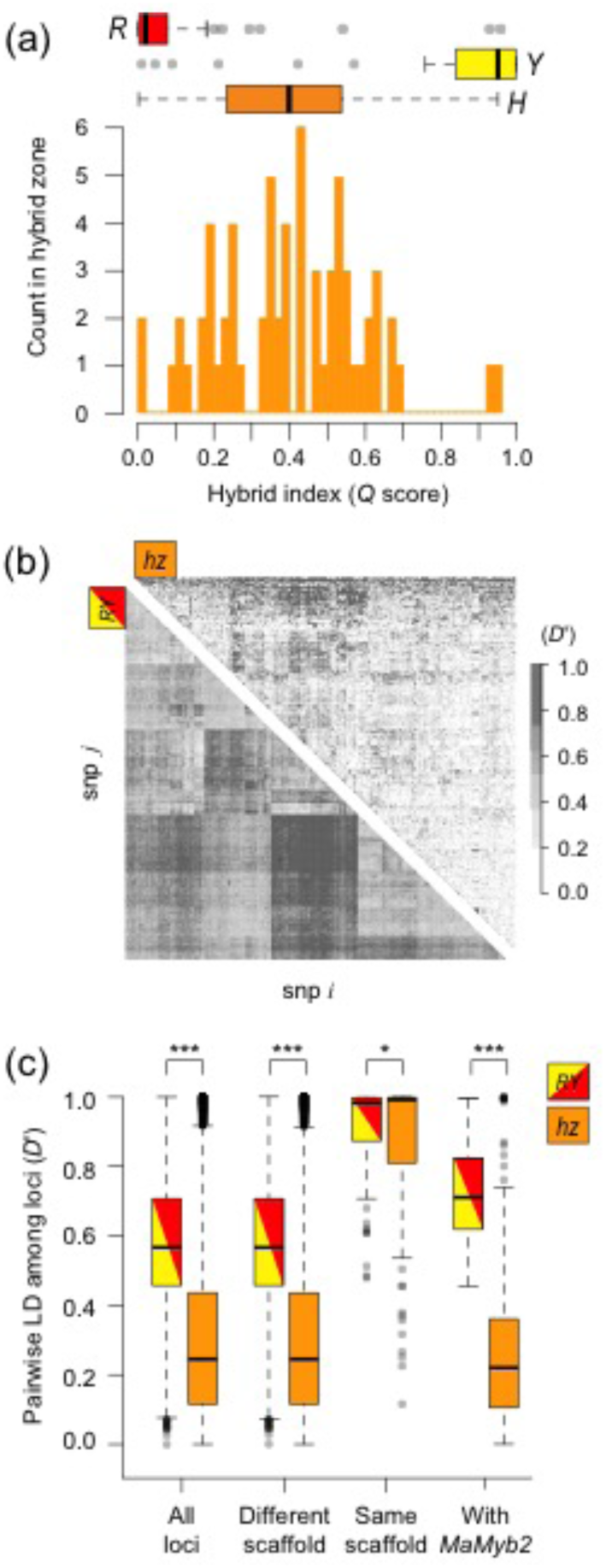
Pattern of admixture and linkage disequilibrium in the hybrid zone. (a) Distributions of hybrid index scores (*Structure Q* score) inside and outside of the hybrid zone. The boxplots show the distributions of *Q* scores for the red (*R*) and yellow (*Y*) ecotypes, and four sample sites inside the hybrid zone (*hz*). The histogram shows a detailed view of the distribution of *Q* scores inside the hybrid zone. (b) Heatmaps of linkage disequilibrium (*D*’) between all pairs of outlier loci (*n* =427) inside the hybrid zone (*hz*), and when the pure red and yellow ecotypes are pooled together (*RY*). Loci are clustered into groups that show elevated linkage disequilibrium with one another. As a consequence, the order of markers differs between the heatmaps. (c) Box plots showing the distribution of linkage disequlibrium (*D*’) for all outlier loci, loci on different genome scaffolds, loci on the same genome scaffold, and for comparisons that include the *MaMyb2-M3* marker. Asterisks indicate the level of significance (based on permutation tests) of mean differences in pairwise *D*’ for comparisons between the hybrid zone and the pooled ‘population’ containing pure red and yellow ecotype individuals (* *p* = 0.02; *** *p* < 9.999 x 10^-5^).

Similarly, we observed significantly reduced linkage disequilibrium (LD) in the hybrid zone relative to the level expected if the ecotypes co-existed without gene flow between them (Fig. 5). Pairwise LD calculated in the hybrid zone was significantly reduced compared to the pooled ‘population’ containing pure red- and yellow-flowered individuals (mean *D*’_RY_ = 0.60, mean *D*’_HZ_ = 0.31; *p* = 9.999 × 10^-5^). In addition, despite large clusters of markers in tight LD in the pooled population, few such clusters were detected in the hybrid zone (Fig. 5b). We also observed significantly reduced LD in the hybrid zone between markers on different genome scaffolds (mean *D*’_RY_ = 0.60, mean *D*’_HZ_ = 0.32; *p* = 9.999 × 10”^5^), as well as for pairwise comparisons that included the *MaMyb2-M3* marker (mean *D*’_RY_ = 0.73, mean *D*’_HZ_ = 0.26; *p* = 9.999 × 10”^5^) (Fig. 5c). In contrast, we observed only marginally lower LD in the hybrid zone between markers on the same genome scaffold, suggesting that LD remained elevated between loci that are less than 74 kb apart (mean *D*’_RY_ = 0.92, mean *D*’_HZ_ = 0.86; *p* = 0.0215).

## Discussion

In this study, we combine an *F_CT_* scan and geographic cline analysis to reveal the genomic signatures of pollinator-mediated divergence between red and yellow ecotypes of *Mimulus aurantiacus*. Overall, our results reveal low genome-wide divergence between the ecotypes, further supporting the conclusion that these taxa are at an early stage of divergence (Sobel and Streisfeld 2015; Stankowski et al. 2015). By contrast, the markers with the steepest clines closely align with the spatial transition in floral traits, suggesting that these loci may reside in or near the genomic regions that contribute to pollinator isolation. Moreover, by taking advantage of the natural hybrid zone between the ecotypes, our data indicate that gene flow and recombination have been extensive, suggesting that the outlier loci are not concentrated in one or a few genomic regions. In addition to elucidating the genomic consequences of pollinator-mediated reproductive isolation in this system, we end by discussing the utility of cline analysis as a spatially explicit framework for future studies of genome-wide variation.

### Genome-wide divergence between the ecotypes

Consistent with a recent origin of the red ecotype from an ancestral yellow-flowered population (Stankowski and Streisfeld 2015), our *F*_CT_ scan revealed very limited genome-wide differentiation between the ecotypes (mean estimate of *F*_CT_ = 0.046). Such low levels of differentiation are predicted in population pairs that are at a very early stage in the speciation process (Feder and Nosil 2012; Nosil et al. 2012; Seehausen et al. 2014). Indeed, our estimate is similar in magnitude to other closely related ecotypes where divergence occurred recently despite gene flow, including apple and hawthorn races of *Rhagoletis pomonella* (*F*_ST_ = 0.035; Feder et al. 2015), wave and crab ecotypes of the intertidal snail *Littorina saxatalis* (*F*_ST_ = 0.027; Butlin et al. 2014) and normal and dwarf forms of the lake whitefish *Coregonus clupeaformis* (*F*_ST_ = 0.046; Herbert et al. 2013).

In addition to revealing the overall level of genome-wide divergence between the ecotypes, our primary goal was to identify loci associated with local adaptation and pollinator isolation in this system. Recent theoretical and empirical studies suggest that that regions of the genome that contribute to local adaptation should show elevated divergence relative to selectively neutral regions (Feder and Nosil 2012; Nosil et al. 2012; Seehausen et al. 2014). However, depending on the strength and timing of selection and the local recombination rate, genome scans based on reduced representation approaches that rely on tight linkage to selected sites may often be underpowered (Arnold et al. 2013). While we cannot rule this possibility out, the presence of a highly differentiated RAD marker (*F*_CT_ =0.66) that is on the same scaffold and ∽ 8 kb away from *MaMyb2*, provides us with confidence that many of the markers in the top 1% of the *F_CT_* distribution are likely to be associated with local adaptation. However, elevated *F_CT_* has a limited capacity for revealing associations between molecular and trait divergence, particularly if phenotypic variation is continuously distributed in space.

As a consequence, we employed cline analysis to provide further support that many of the outlier loci are diverged due to spatially varying selection. While the point estimates of *F*_CT_ for most of these loci are modest, the estimates of the allele frequencies in each tail (*P_min_* and *P_max_*) reveal a complex spatial pattern of allele sharing on both ends of the cline. Almost 80% of the markers are at or near fixation for one allele in one ecotype, while both alleles are present at appreciable frequencies in the other ecotype. This pattern could result from selective sweeps on novel alleles in one of the ecotypes followed by dispersal of that allele to the other ecotype (Pritchard et al. 2010), or from selection on pre-existing, ancestral variation in only one of the ecotypes (Barrett & Schuler 2007). While additional data will be necessary to distinguish between these hypotheses, our analyses provide strong support that selection is responsible for the divergence of these markers despite gene flow.

The estimates of cline center and cline slope provide the most compelling evidence that many of these loci are associated with local adaptation. In a previous study, we showed sharp, coincident clines across the transect for six divergent floral traits (Stankowski et al. 2015). The shape of these clines contrasts with the shallow gradient in genome-wide differentiation, suggesting that the divergent floral traits have been maintained by a common selective agent despite ongoing gene flow, rather than reflecting secondary contact (Streisfeld and Kohn 2005; Stankowski et al. 2015). Thus, we predicted that loci associated with local adaptation should show cline shapes that recapitulate the spatial transition in floral traits. Indeed, we observed 130 RAD markers that have sharp clines and coincide with the narrow phenotypic transition zone between the ecotypes. In ecological models of cline formation and maintenance, the cline center represents the geographic position where the direction of selection switches to favor the alternative form of a trait (i.e., Haldane 1948; Endler 1977; Krukk et al. 1999). Thus, these data are consistent with divergence of these loci due to pollinators, which generally require divergence in multiple traits to maximize attraction and successful pollen transfer (Fenster et al. 2004). Moreover, the clines with the steepest slopes were almost exclusively positioned in this region. Indeed, 56 markers show cline slopes that are steeper than the *MaMyb2-M3* marker. While some may be physically linked to *MaMyb2*, the vast majority of these markers show weak LD with the *MaMyb2-M3* marker in the hybrid zone, suggesting that they are located in different genomic regions. Thus, even though allele frequency differences are generally modest, the relationship between cline center and slope strongly suggests that many of these loci are associated with the primary barrier to gene flow between these ecotypes (Sobel and Streisfeld 2015).

Although the remarkable geographic coincidence of the trait and SNP clines is consistent with local adaptation due to pollinator-mediated selection, there are other explanations for this pattern that must be considered. First, some of these markers could be differentiated due to selective gradients that are unrelated to pollinators but positioned in the same geographic location as the floral traits (Barton and Hewitt 1985). However, there is currently little evidence to support this conclusion. While some vegetative and physiological traits differ between the ecotypes (Hare 2002; Sobel et al. in prep), these traits change in a linear rather than sigmoidal fashion across the study area (Sobel et al. in prep). Thus, we would expect loci associated with adaptive differences in abiotic factors to match the gradual transition in these traits.

In systems where there are genetic incompatibilities between hybridizing taxa, multiple independent clines could become spatially coupled with the cline in phenotypic traits (Bierne et al. 2011). Because endogenous barriers are likely to be attracted to and become trapped by ecological barriers to gene flow, markers contributing to the endogenous barrier need not reside within the genomic regions that contribute to local adaptation (Bierne et al. 2011). However, endogenous barriers to gene flow are effectively absent between the red and yellow ecotypes (Sobel and Streisfeld 2015), suggesting that intrinsic barriers to gene flow are not affecting our ability to identify the loci associated with pollinator isolation. Thus, while additional ecological studies will be necessary to determine whether non-pollinator adaptation is associated with the spatial transition in these markers, current evidence suggests that many of the markers with steep coincident clines likely reside in the genomic regions that underlie the divergent floral traits.

In addition to identifying highly differentiated loci, we also gained insight into how these outliers were distributed throughout the genome. While many studies have shown that outlier loci are scattered throughout the genome (Gompert et al. 2013; Feder et al. 2014; Roesti et al. 2015), others have revealed that they reside in one or a few narrow genomic regions that have diverse phenotypic effects (Lowry and Willis 2010; Fishman et al. 2013; Poelstra et al. 2014). Indeed, the co-localization of adaptive loci appears to have facilitated rapid and robust adaptation in several examples of divergence with gene flow by limiting their breakup in hybrids (Jones et al. 2012; Joron et al. 2011; Lowry and Willis 2010; Twyford and Friedman 2015). At this point, our genome assembly consists of relatively short scaffolds, which limits our ability to establish the physical relationships among the outlier loci in this system. However, our analysis in the hybrid zone suggests that the outlier loci do not co-localize to a small number of genomic regions. Specifically, we observed extensive variation in admixture scores in the hybrid zone, indicating that the divergent alleles are inherited largely independently of one another. In addition, we detected significant reductions in linkage disequilibrium (LD) between pairs of markers in the hybrid zone. Although this could result from the break up of a group of tightly linked loci due to extensive fine-scale recombination in hybrids, our analysis suggests that this is not the case. Indeed, loci on the same genome scaffold (maximum of 74 kb apart) were in strong linkage disequilibrium both inside and outside of the hybrid zone. Thus, our results suggest that many of the outlier loci reside either on different chromosomes or in tightly linked co-linear regions of the same chromosome. This result is in agreement with the substantial breakup of divergent floral traits in these same hybrid populations, and in an experimental F_2_ population where selection was relaxed (Stankowski et al. 2015). Future studies that take advantage of chromosome-length genomic scaffolds will help to resolve the full extent of LD across the genome to determine the potential for structural variants that might limit recombination among loci.

### Geographic cline analysis as a tool for studying genome-wide divergence

In addition to characterizing the genomics of floral divergence between the red and yellow ecotypes of *M. aurantiacus*, an additional goal of our study was to explore the utility of geographic cline analysis as a tool for studying genome-wide patterns of variation. Geographic cline analysis has long been used for studying barriers to gene flow between closely related taxa. However, despite a call for better integration of cline theory into genomic studies of adaptive divergence and speciation (Bierne et al. 2011), geographic cline analysis has not been applied to large datasets with the explicit purpose of studying patterns of genome-wide variation. This seems to reflect the history of development and use of cline analysis, and the relative difficulty of applying cline analysis to large marker datasets.

While clines have long been recognized as excellent systems for developing and testing ideas about speciation (Huxley 1938; Haldane 1948; Bazykin 1969; Clarke 1966; Endler 1977), the majority of cline theory was developed in the 1980s and 90s to provide a powerful framework for making detailed inferences about the nature and strength of barriers to gene flow between hybridizing taxa (Barton 1983; Barton & Hewitt 1985; Szymura and Barton 1986; Mallet and Barton 1989; Barton and Gale 1993). Although this method requires just a handful of differentially fixed, unlinked loci, cline analysis can be applied to larger datasets, with the goal of studying patterns of genome-wide variation. For example, consider a scenario where local adaptation across a sharp ecological gradient arises from selection on standing genetic variation. In this case, considerable allele sharing between divergent populations is expected at sites linked to the causal variants (Hermisson and Pennings 2005; Pritchard et al. 2010). Although allele frequency differences at linked sites may be relatively small, any difference in allele frequency between populations should manifest itself as a sharp cline due to the indirect effects of selection on linked variants. Indeed, a model of ecological cline maintenance for a selected locus and a tightly linked neutral locus predicts the formation of clines with similar shapes, though the level of allele sharing is higher at the neutral marker (Durrett et al. 2000). Similarly, in polygenic models of adaptation, where traits are controlled by many loci each of small effect, smaller differences in allele frequency are expected among diverging populations even at causal loci (Pritchard et al. 2010).

In our study, the loci with cline shapes that recapitulated the spatial patterns of floral trait variation were associated with modest allele frequency differences. In contrast, only 106 of the 427 outlier loci showed an allele frequency difference across the transect greater than 0.8 (Δ*P* > 0.8). Even more striking, these markers tended to have cline shapes that were neither coincident nor concordant with the clines in floral traits. Rather, they tended to show very broad clines that were often positioned large distances from the transition in the floral traits. While these loci may be associated with other forms of local adaptation or reflect a potentially complex history of divergence, our analysis suggests that they do not make a major contribution to floral trait divergence. Thus, if we had limited our cline analysis only to these loci, we would draw very different conclusions about the pattern of genome-wide divergence in this system

Another reason that cline analysis has not been applied to genome-wide data is that it is more complex and computationally intensive compared with other methods. For example, *F_ST_* and other similar measures of differentiation are easily calculated, even for thousands to millions of loci. Genomic cline analyses, which fit functions to allele frequency data plotted against a hybrid index instead of geographic distance, are fully automated (i.e., Gompert and Buerkle 2012). However, under certain situations, geographic cline analysis has several advantages over these methods. For example, neither *F_ST_* nor genomic cline analyses can incorporate spatial, phenotypic, or environmental data into studies of genome-wide variation. Moreover, our analysis revealed that key cline parameters, including the cline center and slope, showed only weak correlations with *F*_CT_. These results indicate that cline analysis provides a more detailed view of divergence in this system that can be related directly to the patterns of divergence in ecologically important traits.

As a preliminary study of clinal variation in this system, we only fitted clines to the top 1% of the *F*_CT_ distribution, but we argue that full genome-wide cline analysis at all variable sites can be conducted in conjunction with traditional genome scans. Efficient software is now available to allow the automated fitting of cline models to large datasets (Derryberry et al. 2015). Ideally, the fit of a cline model would be compared to the fit of a null model (slope = 0) to identify loci that show clinal variation. The resulting cline parameters then can be mapped across chromosomes to reveal the consequences of selection and reproductive isolation across the entire genome, as has been done in genomes scans using other population genetic statistics (i.e., Hohenlohe et al. 2010 Burri et al. 2015). While the small scaffolds in our current genome assembly preclude this analysis, our analysis of SNPs from the same genomic scaffold indicate that cline shape parameters are correlated across small chromosomal regions, demonstrating the potential of this method in our system. The interpretation of genome-wide patterns of clinal variation will be aided by genomic models of cline formation and maintenance across a range of divergence histories. Integration of these genome-wide patterns of divergence with studies of QTL mapping of trait variation, historical demographic modeling and haplotype-based analyses will enhance our understanding of ecological divergence in this system, and other examples of divergence with gene flow.

## Acknowledgments

We would like to thank Susie Bassham for advice on sequencing library preparation, Josh Burkhart for assistance with the assembly, and Julian Catchen for modifying the *Stacks* pipeline. Madeline Chase, Thomas Nelson and William Cresko provided fruitful discussion. Lorne Curran provided computer support. The project was supported by National Science Foundation grant: DEB-1258199.

## Data Archiving

Sequence data will been submitted to the Short Read Archive

